# Proteomic and Kinetic Characterization of Prion Seeding in Distinct Human CJD Strains Unveils Early Diagnostic Biomarkers

**DOI:** 10.1101/2025.09.17.676758

**Authors:** Saima Zafar, Neelam Younas, Jean-Yves Douet, Mathias Schmitz, Niccolò Candelise, Susana Correia, Olivier Andréoletti, Inga Zerr

## Abstract

To enhance understanding of early diagnosis, differential disease progression rates, and the molecular profiles of human prion strains, we analyzed prion seeding activity over time in Creutzfeldt-Jakob disease (CJD)-infected mice using the real-time quaking-induced conversion (RT-QuIC) assay. Our previous work highlighted pre- clinical alterations in endocytic machinery and cytoskeleton-associated responses in CJD-affected brain regions. In this study, infectious prion strains derived from human CJD-MM1 and VV2 brain tissues were inoculated into tg340, tg361 (expressing approximately four times the human PrP-M129 and PrP-V129, respectively), and tg650 (expressing approximately six times the human PrP-M129) mice. A total of 188 brain samples (cortex) were analyzed from confirmed CJD-infected mice and control mice at both pre-clinical and clinical stages of the disease. Notably, we observed region-specific and PrP strain-specific differences in seeding activity at the pre-clinical stage of disease in CJD-MM1 and VV2 infected mice. The lag phase between the positive response ranged from 7.5 to 24.5 hours across all regions and disease stages. In the cortex, CJD-MM1-infected tg340 mice exhibited a prolonged lag phase (∼24.5 hours), while CJD-VV2-infected tg341 mice showed minimal seeding response and relative fluorescence unit (RFU) signal rates. In contrast, the cerebellum of VV2 clinical stage mice exhibited a shorter lag phase, and VV2 preclinical stage mice showed significantly lower RFU signal rates.

Proteomic profiling via SWATH-MS identified 500 and 682 differentially expressed proteins in the MM1 and VV2 models, respectively. Key proteins such as Gnl1, Stxbp1, Pllp, Gps1, Nefh, Ahsa1, Rala, Cacybp, Pdk1, Unc13a, Rab21, Rraga, Ppp1r9a, Eif4b, Atp2b2, Vps51, H2afx in CJD-MM1 and Gm10358, Tnc, Calb2, Ppm1h, Dnaja1, Gm45808, Hpcal1, Prkca, Dock3, Syn2, Agap2, Tmem126a, Fdps, Ndufa4 in CJD- VV2 showed significant alterations at the early pre-clinical stages, correlating with detectable prion replication. These molecular shifts highlight potential early-stage diagnostic biomarkers. Functional analyses revealed that both MM1 and VV2 subtypes engage early compensatory responses; however, MM1 primarily involves metabolic reprogramming and enhanced vesicle clearance, while VV2 is characterized by pronounced disturbances in calcium signaling and structural integrity at early stage of the disease. These findings emphasize the utility of RT-QuIC and proteomics in characterizing prion seeding and progression, providing valuable insights into the molecular mechanisms underlying prion diseases and potential early diagnostic markers.

## 1. Introduction

Creutzfeldt-Jakob disease (CJD), a human prion disease, is characterized by the accumulation of misfolded prion proteins (PrP^Sc^), which propagate infectious forms of prion protein, leading to neurodegeneration. The disease is classified into different subtypes based on the polymorphism at codon 129 of the prion protein gene (PRNP) and the electrophoretic profile of the PrP^Sc^ isoform. The MM1 and VV2 subtypes are the most common human prion strains and have been shown to exhibit distinct patterns of disease progression, neuropathological features, and molecular alterations (Peden AH 2004, Collinge J 2006). More recent studies have emphasized the variability in prion strain behavior across species and host-genotype backgrounds, highlighting the complexity of prion disease pathogenesis (Prusiner SB 2015, Schmitz M 2016).

Early diagnosis and understanding of prion diseases have proven difficult due to the lack of clear diagnostic markers in the preclinical phase. While clinical diagnosis often occurs at a late stage of disease, the early molecular changes associated with prion propagation remain poorly understood (Altuna, et al. 2022, Wang 2019). Recent studies have highlighted that prion seeding activity in tissues, such as the brain, can be detected in preclinical stages of disease using real-time quaking-induced conversion (RT-QuIC) assays, which have become an important tool for monitoring prion infection dynamics (Atarashi R 2011, Schmitz M, 2016, Zerr I, 2020). Moreover, novel biomarkers for prion diseases have been identified using liquid biopsy techniques, allowing for the detection of prion seeding in blood and cerebrospinal fluid (CSF) (Pérez-Lázaro S 2024, Vallabh SM 2024, Küçükali F 2025).

To address this gap, our study investigates the early preclinical seeding activity and molecular profiling in CJD-infected transgenic mice expressing human PrP-M129 or PrP-V129. These mice, tg340 (PrPM-129) and tg361 (PrP-V129), have been shown to replicate prion strains in a manner that is consistent with human disease progression. By inoculating these transgenic mice with MM1 and VV2 prion strains and monitoring the seeding activity over time using RT-QuIC, we were able to track disease progression at multiple stages, ranging from the early preclinical to late symptomatic phases.

Furthermore, we performed a proteomic analysis using Sequential Windowed Acquisition of All Theoretical Fragment Ion Mass Spectra (SWATH-MS) to identify differentially expressed proteins at early stages of prion infection (Fig.1b). This approach provides an unbiased, comprehensive profile of molecular changes occurring during prion propagation and has been successfully used to identify potential biomarkers for prion diseases (Shafiq M 2021).

Our findings indicate that early proteomic alterations in MM1 and VV2 infected mice correlate with prion replication dynamics, offering new insights into the molecular underpinnings of prion diseases and potential targets for early diagnostic biomarkers. We report for the first time that differential seeding activity and molecular changes associated with prion infection are detectable as early as the preclinical stage. These findings underscore the potential of RT-QuIC and proteomic profiling to serve as valuable tools for understanding prion disease progression and identifying early diagnostic markers in human prion diseases.

## 2. Materials and Methods

### 2.1 Ethics statement

All animal experiments were conducted in accordance with the ethical standards established by the Regierungspräsidium Tübingen (Regional Council), under experimental approval number FLI 231/07 (file reference 35/9185.81-2). Additionally, all procedures complied with institutional and French national guidelines, in line with the European Community Council Directive 86/609/EEC. The experimental protocol was approved by the INRA Toulouse/ENVT Ethics Committee

### 2.2 sCJD MM1 and VV2 transgenic mouse lines

Both transgenic mouse lines, tg340(tgMet) and tg361(tgVal), that express human PrP methionine at codon 129 or valine at codon 129, respectively, in a PrP knockout background were used, as described previously (Zafar S, 2015, Zafar S, 2017). Both mouse lines are homozygous for the human PRNP and overexpress about 4-fold level of human PrP^C^. Inocula were prepared from sCJD-MM1 and VV2 brain tissues as 10% (w/v) homogenates. Six- to ten-week-old mice were anesthetized and inoculated with 2 mg of brain homogenate in the right parietal lobe using a 25-gauge disposable hypodermic needle. Mice were observed on a daily basis, the neurological assessment was performed weekly. Animals were sacrificed when disease progression was evident, or at the end of lifespan. Animals were euthanized, and the brain was collected at 4 time points [60-, 120-, 160- and 180-days post inoculation (dpi)] corresponding to early pre-symptomatic, late pre-symptomatic, early symptomatic and late symptomatic stages, respectively. All mice (n=six animals per group and per time point) were anesthetized, decapitated, and brain was harvested.

Brain tissue was frozen at −80 °C for subsequent protein extraction. For each isolate, survival time was calculated as the mean number of days post-inoculation (dpi) among mice that tested positive for PrP^Sc^. The infection rate was defined as the percentage of PrP^Sc^-positive mice out of the total number of inoculated animals.

### 2.3 Preparation of brain tissue homogenate for proteomics analysis

Frontal cortex tissues from spAD, rpAD, sCJD (MM1, VV2 subtypes), and non-demented control subjects, and cortical tissues from mouse brain were homogenized as previously described (Llorens F 2014, Zafar S 2015). Briefly, brain tissues were lysed (10%, wt/vol) using tissue lyser LT (Qiagen/Hilden, Germany) in ice-cold tissue lysis buffer supplemented with protease and phosphatase inhibitors (Roche, Germany). The lysates were centrifuged at 14000 rpm for 30 min at 4 °C to remove insoluble debris. Protein concentration in the supernatants was determined by the Bradford method, using bovine serum albumin (BSA) as a standard. Bradford assay dye reagent was diluted with ddH_2_0 at a ratio of 1:5. Protein standard (BSA) dilutions were prepared ranging from 0.0-1.0 mg/ ml in ddH20. Total Protein extracts from experimental samples were diluted at a ratio of 1:20. From this dilution, 20 μl was mixed with 980 μl of Bradford working solution, and the mixture was incubated for 10 min. at RT. Optical density (OD) of the different dilutions of protein standard and experimental samples was measured at a wavelength of 595 nm. Protein concentration was calculated with the help of OD values by using Microsoft Office 2013 Excel software.

### 2.4 SWATH-MS (Sequential Windowed Acquisition of All Theoretical Fragment Ion Mass Spectra) global proteomics

#### 2.4.1 Library preparation

All samples were lysed as above to isolate total protein extracts and protein concentration was measured by Bradford assay. We prepared the spectral peptide library as mentioned previously (Shafiq M 2021). Briefly, Equal amounts of protein from each sample were pooled and fractionated into eight parts using a high-pH reverse-phase spin column (Thermo Fisher Scientific) to generate a spectral peptide library. Digested peptides (1.5 µg per run) were analyzed on an Eksigent nanoLC425 system coupled to a TripleTOF 5600+ mass spectrometer with Nanospray III ion source. Peptides were separated on a C18 column using a 100-minute gradient and analyzed using a Top30 data-dependent acquisition method. MS1 scans (m/z 380–1250) and MS2 scans (m/z 180–1500) were collected at resolutions of 35,000 and 17,500 FWHM, respectively. CID was used for fragmentation, and dynamic exclusion was set to 15 s. Each fraction was analyzed in duplicate for spectral library construction.

#### 2.4.2 Quantitative SWATH measurement

For quantitative SWATH measurement, as mentioned previously by Shafiq *et al.,* 2021, MS/MS data were acquired using 100 variable windows across a 400–1200 m/z range, with fragments collected from 180–1500 m/z for 40 ms per segment, and a 250 ms survey scan, yielding a 4.3 s cycle time. Two technical replicates were acquired per biological sample. Protein identification was performed using ProteinPilot v5.0 (AB Sciex) with “thorough” settings, searching against the UniProtKB human proteome (02/2017, 92,928 entries) plus 51 common contaminants, identifying 4,537 proteins at 1% FDR. SWATH peak extraction and spectral library generation were conducted in PeakView v2.1 using the SWATH microApp v2.0, with retention time correction and peak area extraction performed at 1% FDR (Gillet, et al. 2012). The resulting areas of peaks were then summed up to peptide and finally corresponding protein area values, which were proceeded for further statistical analysis.

### 2.5 Differential expression analysis of proteomic alterations

To identify proteins that were differentially expressed at different stages of the disease in comparison to control, t-test (two sided) was performed using the Perseus software platform (Version 1.5.0.31) (Zafar, et al. 2022), with a significance cut-off of p < 0.05. Given the exploratory nature of this discovery-based proteomic workflow post-hoc correction was not applied in the current data analysis. Volcano plots were calculated by Perseus software in which the fold change (FC) was transformed using the log2 function, so that the data is centered on zero, while the p-values were −log10 transformed. Volcano plots were prepared in R (version 2.8.0) followed by editing with Inkscape. Fold Change (FC) for all comparison’s threshold at ±1.5 with a p-value less than 0.05 as a cutoff for significance. The proteins with significant differential expression are indicated with red (up-regulated) and green (down-regulated) in the volcano plots. Significantly proteins were used to make heatmap by Perseus Software (Version 1.5.0.31).

### 2.6 Gene Ontology (GO) analysis and functional network mapping

To gain functional insights from proteomics data, three enrichment approaches were applied. GO term enrichment (biological process, molecular function, and cellular component) was first assessed using Fisher’s exact test in Perseus. GO Slim summaries were then obtained via overrepresentation analysis using WebGestalt. Finally, Ingenuity Pathway Analysis (IPA) was conducted to identify canonical pathways linked to Tau-related protein changes.

Predicted target genes for each protein were analyzed using the ToppFun module of the ToppGene Suite (http://toppgene.cchmc.org/), focusing on Gene Ontology (GO) terms for molecular function, biological process, and cellular component. Enrichment significance was assessed using Bonferroni correction with a p-value threshold of 0.05.

### 2.7 RT-QuIC analysis

RT-QuIC was performed as previously described (Schmitz M, 2016). Reactions (100 µL total volume) included 85 µL of buffer containing 5× PBS (pH 6.9), 170 mM NaCl, 1 mM EDTA, 10 µM Thioflavin-T, and 0.1 mg/mL recombinant PrP^C^, seeded with 15 µL of 10% brain homogenate (10⁻³ dilution). Samples were run in triplicate in 96-well black optical plates, sealed, and incubated at 42 °C for 80 hours in a FLUO Star OPTIMA reader with alternating 1-minute shaking (600 rpm) and 1-minute rest cycles. Thioflavin-T fluorescence (excitation 450 nm, emission 480 nm) was recorded every 30 minutes to monitor β-sheet formation.

### 2.8 Statistical analysis

All data in the study was obtained from at least three independent experiments. All results are expressed as mean ± SEM. Densitometric analysis of 1-DE gels was performed using Lab image (version 2.7.1 Kapelan, Leipzig, Germany) software. Statistical analysis was performed with GraphPad Prism 6.01. Proteomics data was analyzed using Perseus software platform with adjustments for multiple testing. Hierarchical clustering analysis was also performed with Perseus software. Two-dimensional interaction plots were plotted in R (pre-release version 2.8.0) followed by editing with Inkscape. Student’s *t*-test was used for comparisons between two groups. One-way ANOVA followed by *Tuckey post-hoc* test was used for comparisons between three or more groups. Statistical significance was considered for p-value <0.05.

## 3. Results

### 3.1 Early preclinical seeding activity of prion protein during prion infection detected by RT-QuIC analysis

To determine the timing of initial prion seeding during infection, we investigated prion protein seeding activity over time using RT-QuIC. Humanized mice of the appropriate genetic background were intracerebrally inoculated with brain homogenates from human sCJD-MM1, sCJD-VV2 cases, or non-infectious controls (Fig. 1a). Brain samples from the cortex region were collected at four defined time points—60, 120, 160, and 180 days post-inoculation (dpi)—corresponding to early preclinical, late preclinical, early clinical, and late clinical stages, respectively. A total of 96 samples were analyzed, with six biological replicates per time point for both prion-infected and control groups. RT-QuIC was then performed on these samples to assess the progression of seeding activity during disease development.

**Fig. 1:**
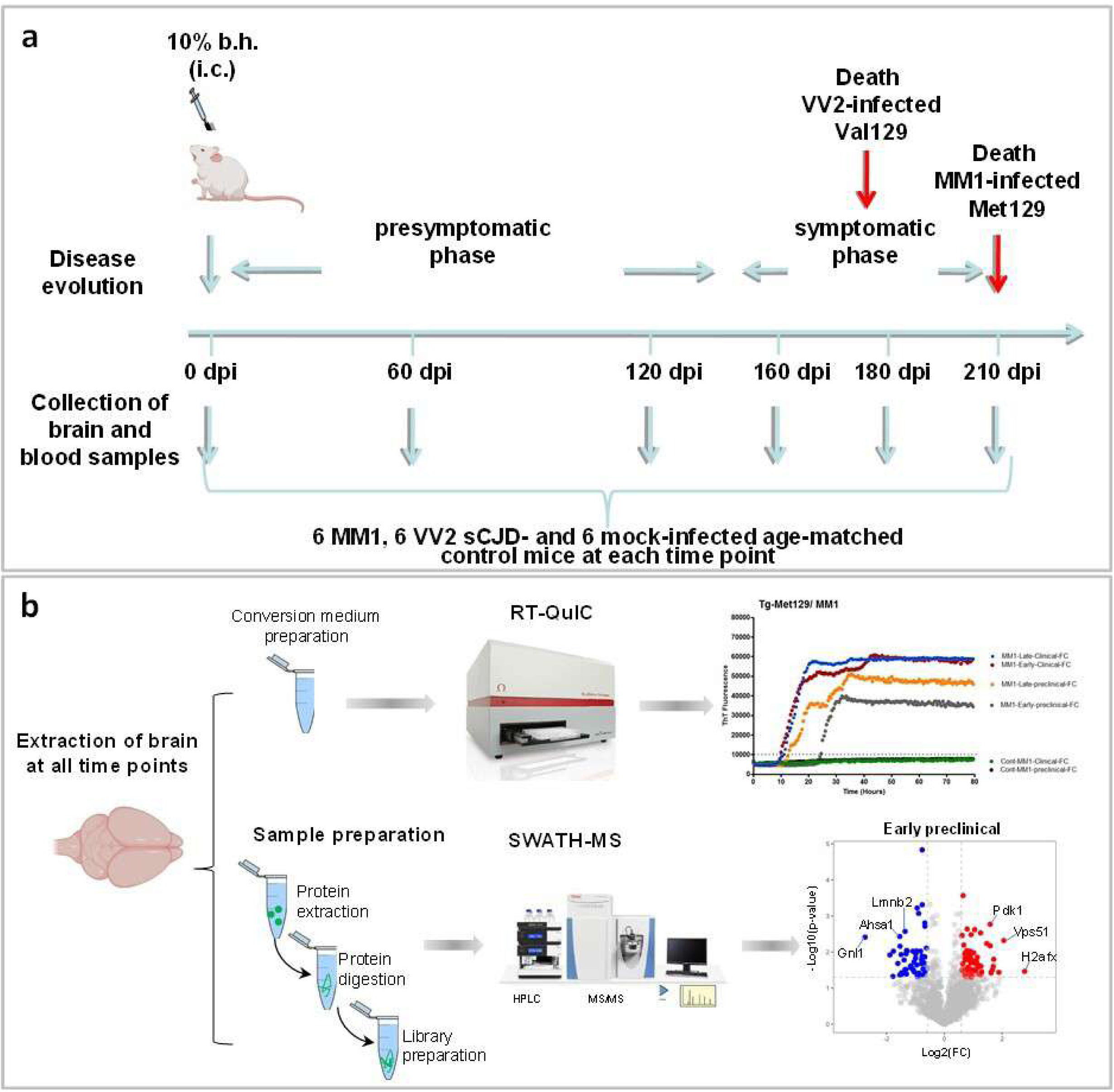
Identification of prion seeding and proteomic alterations during prion disease progression. **a)** Timeline of prion inoculation and sample collection at preclinical and clinical stages of the disease. Brain homogenates (b.h.) (10%) from sporadic CJD-MM1 or VV2 patients were inoculated into humanized mouse models. **b)** Workflow of seeding and proteomics analysis. To find out the time of initial prion seeding during prion infection, the seeding activity of prion protein was investigated in a time-dependent manner using RT-QuIC (Real time quacking induced conversion), upper part of the figure. Proteomic alterations were measured using quantitative mass spectrometry method known as SWATH-MS (Sequential Window Acquisition of All Theoretical Mass Spectra). Volcano plots are showing p-values (-log10, y-axis) and the fold change (log_2_FC, x-axis) at 60 days post inoculation: dpi. The data points above the grey dashed lines are representing proteins with a p-value < 0.05 and FC > ±1.5, as significant candidate hits. Red (upregulated), blue (down regulated).

Strikingly, prion seeding activity was detectable as early as the preclinical stage. Positive RT-QuIC signals were observed at 60 dpi in both sCJD-MM1 and sCJD-VV2 inoculated mouse models (Fig. 2a, b). In contrast, no seeding activity was detected at any stage in mice inoculated with non-infectious brain homogenates (controls) (Fig. 2a, b). In MM1-inoculated mice, RT-QuIC reactivity progressively increased from early preclinical to late clinical stages (Fig. 2a). Conversely, in sCJD-VV2 inoculated mice, reactivity rose from early preclinical to early clinical stages but declined at the late clinical stage (Fig. 2b).

**Fig. 2:**
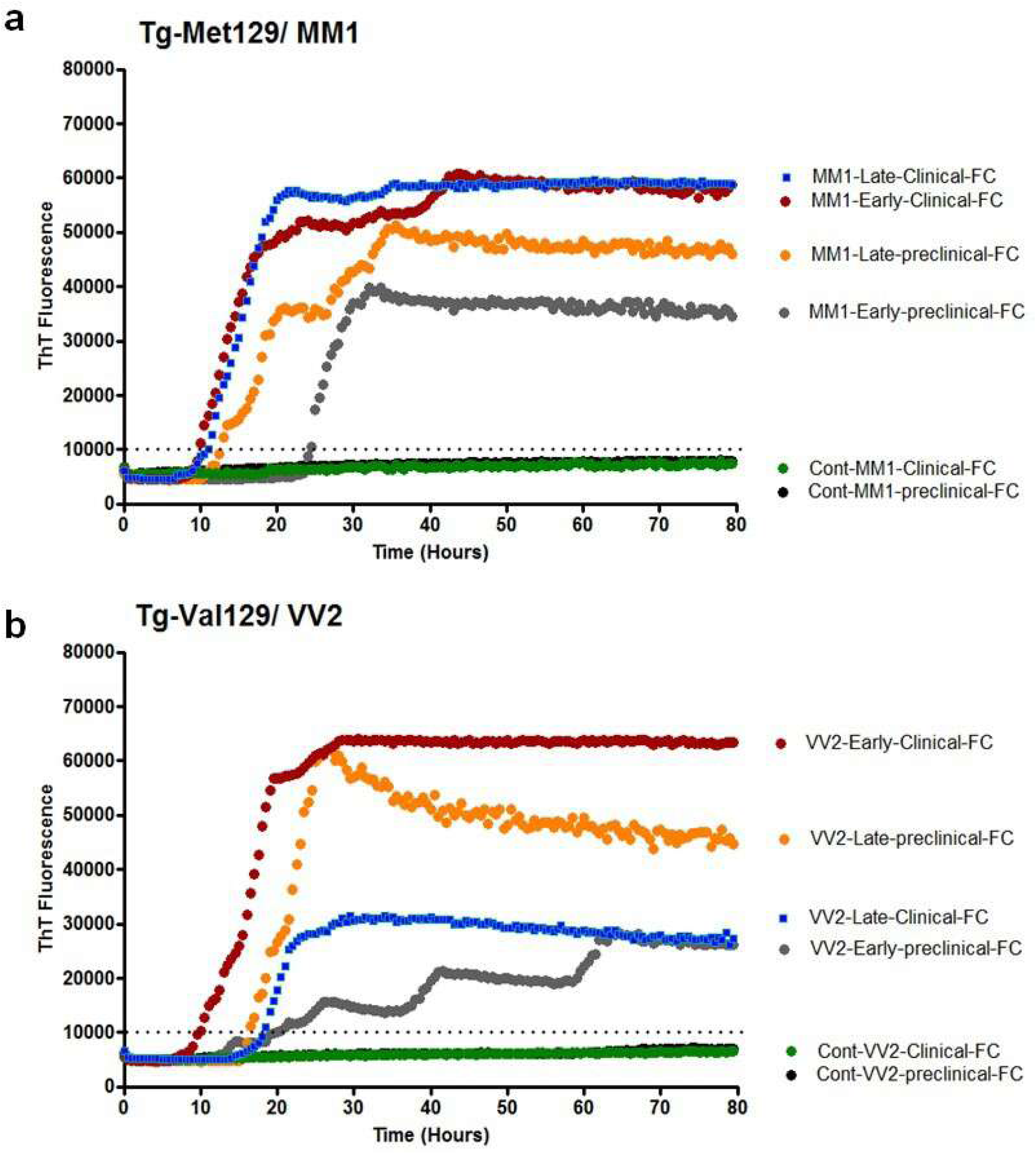
RT-QuIC detection of prion seeding during prion infection. The prion seeding activity was investigated in **a)** CJD-MM1- and **b**) CJD-VV2-inoculated mice and controls at four time points corresponding to early preclinical (60dpi, grey line), late preclinical (120 dpi, orange line), early clinical (160 doi, red line) and late clinical (180 dpi, blue line). Tissue homogenates from control animals (black: preclinical, green: clinical) showed no response. To seed RT-QuIC reactions, 10^−3^ brain tissue dilutions were used. ThT fluorescence was measured at one-hour intervals and average values from quadruplicate wells were plotted as a function of time. The fluorescence threshold was 10,000 rfu, which was the basis of determining positive RT-QuIC response. Dpi: dys post inoculation.

A fluorescence threshold of 10,000 rfu was set to define a positive RT-QuIC response. The lag phase—the time required to cross this threshold—varied between 7.5 and 24.5 hours across different disease stages. At the early preclinical stage, CJD-MM1 infected mice exhibited a longer lag phase (∼24.5 hours) compared to CJD-VV2 inoculated mice (∼15 hours) (Fig. 2a, b). In CJD-VV2 mice, the shortest lag phase (∼7.5 hours) was observed at the early clinical stage, while other stages showed lag phases around 18 hours (Fig. 2b).

Overall, we observed distinct seeding kinetics during prion infection, with prion seeding detectable as early as the preclinical phase. Responses surpassing the threshold typically appeared within 0–10 hours post-assay initiation for both mouse models, albeit at different disease stages. Late clinical stage isolates from CJD-MM1 mice showed positive signals within 0–10 hours (Fig. 3a), whereas positive responses in CJD-VV2 mice were detected as early as the clinical stage (Fig. 3b). Between 0–20 hours, positive responses were observed at late preclinical, early clinical, and late clinical stages for both models, with varying signal intensities (Fig. 3c, d). Notably, early preclinical CJD-VV2 samples reached the threshold after approximately 20 hours. Extending the observation window to 0–48 hours revealed positive signals across all preclinical and clinical stages (Fig. 3e, f). No responses exceeding the threshold were detected in control mice at any time point (Fig. 3a–f).

**Fig. 3:**
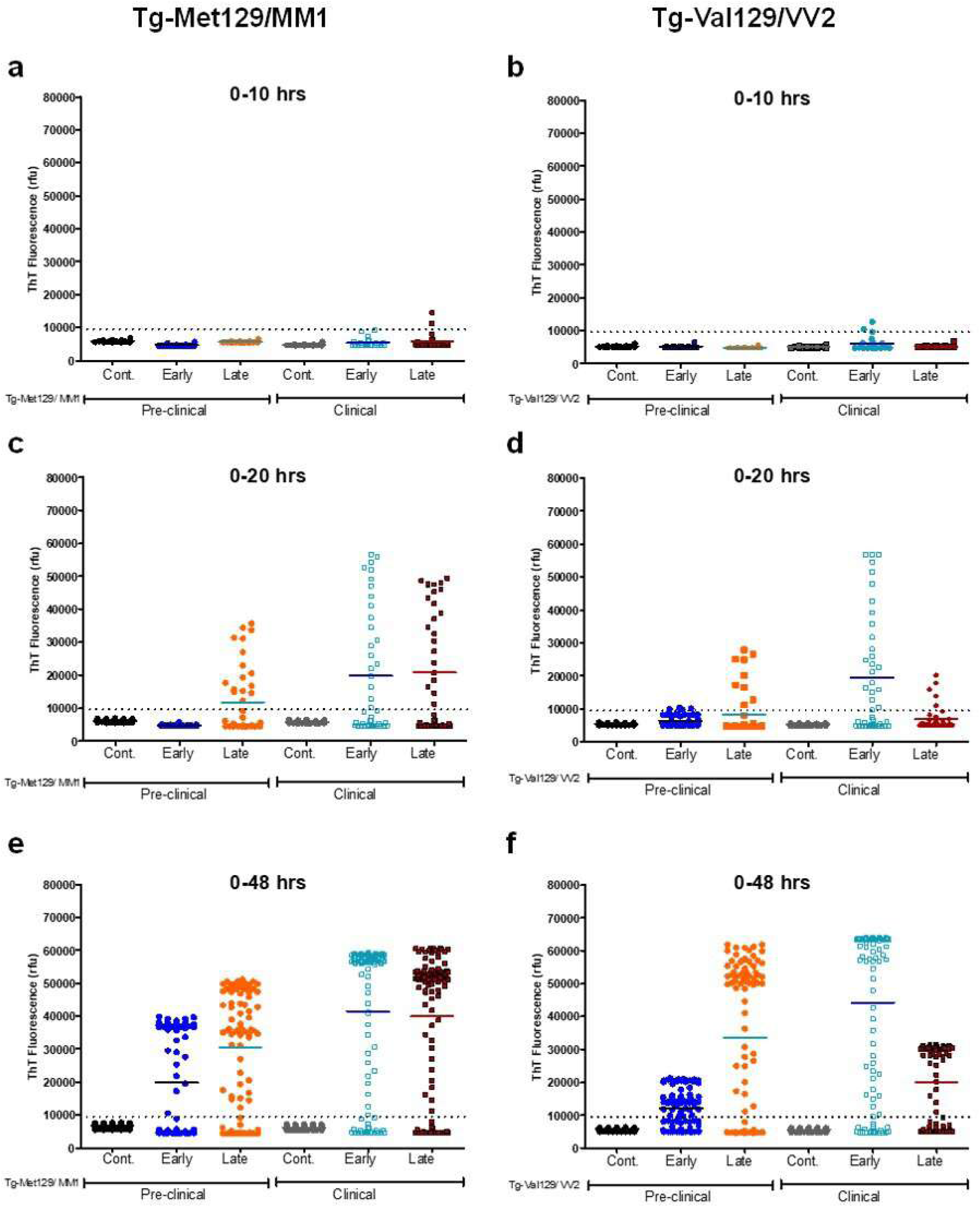
Timing of initial seeding of prion protein during disease progression via RT-QuIC. The prion seeding activity was investigated in (**a, c, e)** CJD-MM1- and (**b, d, f**) CJD-VV2-inoculated mice and controls (black and blue dots) at four time points corresponding to early preclinical (60dpi, dark blue), late preclinical (120 dpi, orange), early clinical (160 dpi, light blue) and late clinical (180 dpi, red) time points. Y axis is showing THT fluorescence (rfu) and x-axis is showing time points. Each sample was tested in quadruplicates, and each plot corresponds to one mouse (n = 6). RFU: relative fluorescence units, dpi: days post inoculation.

### 3.2 Preclinical proteomics alterations in CJD-MM1 and -VV2 mice

Next, to identify molecular changes associated with initial prion seeding events and how they change during the progression of the disease, we performed time-dependent quantitative proteomics analysis. Total protein isolates from cortex of inoculated CJD-MM1 and VV2 mice and mock-infected mice (control) were extracted and subjected to SWATH-MS analysis (n=48, 3 biological replicates per time point from both prion-infected and control mice). There were 1601 proteins quantified at critical FDR of 1%. Firstly, to identify differentially expressed proteins (DEPs) between inoculated and control mice, proteins were uploaded to Perseus software, followed by Welsch’s t-test. This analysis revealed 500 proteins that were differentially expressed between sCJD-MM1 and control mice (**p_value < 0.05; Fig. 4 a-d, S1 Table**). Additionally, a total of 682 proteins were differentially expressed between sCJD-VV2 and control mice (**p_value < 0.05; Fig 4 e-h, S2 Table**).

**Figure 4:**
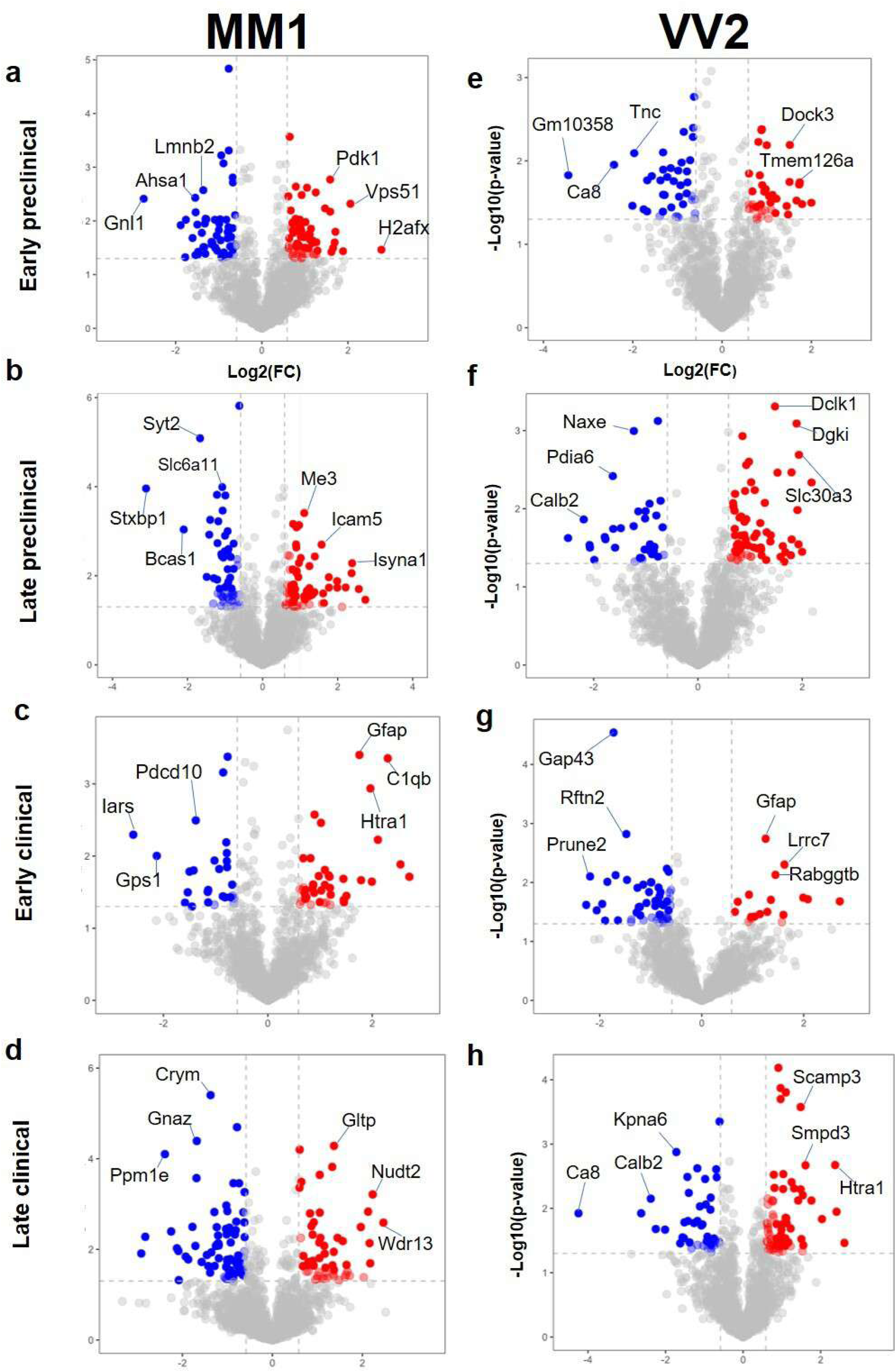
Temporal alterations at proteome level during prion disease progression: The proteomic alterations were investigated in **a-d)** CJD-MM1- and **e-h)** CJD-VV2-inoculated mice and controls at four time points corresponding to early preclinical (60 dpi), late preclinical (120 dpi), early clinical (160 dpi) and late clinical (180 dpi). Volcano plots (Welch’s t-test) are presenting the differentially expressed proteins, p-values (-log_10_ at y-axis) and the fold change (log_2_FC at x-axis). The data points (proteins) above the grey dashed lines are representing significant hits with a p-value < 0.05 and FC > ±1.5, upregulated (red) and downregulated (blue). Dpi: days post inoculation.

Proteomic alterations were already detectable at the early preclinical stage, with significant changes observed (fold change > ±1.5, p < 0.05) (Fig. 4a–h). Notable early changes included proteins such as Pdk1, Vps51, Lmnb2, and Ahsa1 in CJD-MM1 mice, and Dock3, Tnc, Ca8, and Gm10358 in CJD-VV2 mice. To examine temporal dynamics, significantly regulated proteins were subjected to hierarchical clustering to generate a heatmap, where columns represent disease stages (preclinical and clinical) and rows indicate individual proteins. This analysis revealed three distinct expression patterns each for CJD-MM1 and CJD-VV2 (Fig. 5a, b).

**Figure 5:**
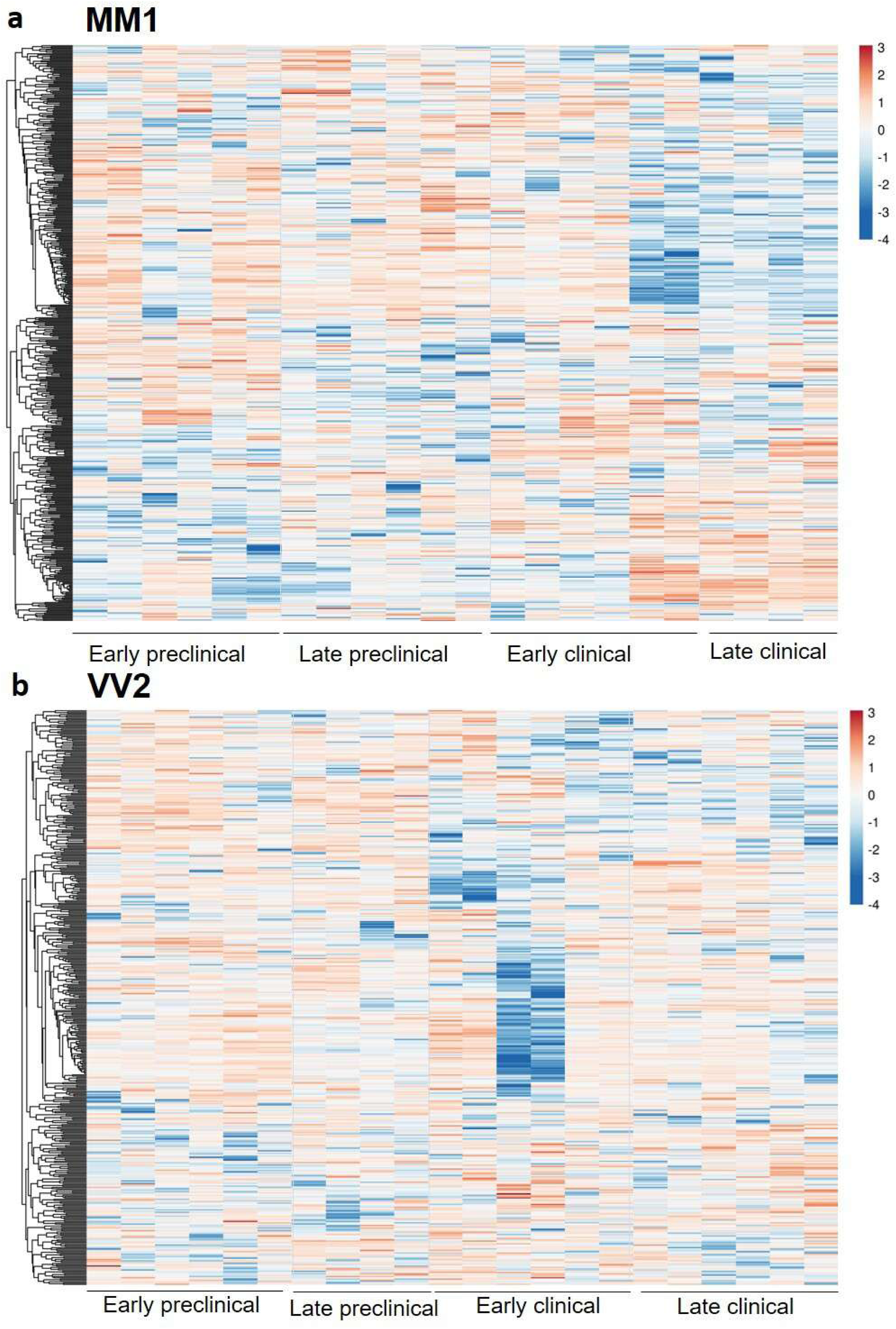
Temporal pattern of proteomic alterations in CJD-MM1- and CJD-VV2- inoculated mice. The heat map is showing pattern of proteomic alterations over the course of the disease. The log_2_-transformed expression values were Z-score normalized for every biological replicate. Columns of the heat map are showing biological replicates at four time points (early preclinical, late preclinical, early clinical and late clinical) and the rows are indicating proteins.

Strikingly, proteomic changes exhibited a clear temporal pattern. Proteins altered at preclinical stages were largely distinct from those dysregulated at clinical stages in both CJD variants (Fig. 5; Supplementary Tables S1 and S2). Importantly, these early proteomic changes (starting at 60 dpi) coincided with the onset of detectable prion seeding activity identified by RT-QuIC (Fig. 2), suggesting that early proteomic alterations reflect a molecular response to initial prion replication and track with disease progression.

Furthermore, we performed comparative Venn diagram analyses (Figure 6). Each panel (a–d) in Figure 6 illustrates the overlapping of proteins identified in MM1 and VV2-infected mice at 60, 120, 160 and 180 dpi. Panel a compares from MM1-infected samples at 60, 120, 160 and 180 dpi. A shared set of 5 proteins (Camkv, Cntnap1, Epb41I3, Stxbp1, Sirt2) were found across all time points, while each time point also exhibited a substantial number of unique proteins, with MM1-180 displaying the highest number of unique modified 126 proteins. Panel b examines proteins from VV2-infected samples across the same time points (60, 120, 160, 180 dpi). Only 1 protein (Basp1) was consistently expressed across all four time points, with VV2-180 showing the greatest number of unique proteins (130). Notably, the overlap between VV2-120 and VV2-160 was limited to just 3 proteins, suggesting highly dynamic transcriptomic changes. Fig. 6 Panel c presents a cross-strain and cross-time comparison involving MM1 and VV2 at 60 and 120 dpi. Only 2 proteins (Plp1, Slc4a4) were found to be common across all four conditions, while MM1-dpi60 and VV2-dpi60 shared 8 proteins (Slc44a1, Pgm1, Cnp, Endod1, Tpm3, Kif2a, Agap2, Sirt2) independently, and each condition maintained a considerable set of unique proteins. Panel d focuses on a comparison among VV2-160, VV2-180, MM1-160, and MM1-180. Only 7 proteins (Basp1, Crym, Mog, Ywhaq, Map2k1, Tubb2a, Tpi1) were shared among three conditions (excluding VV2-160), with no proteins shared across all four. MM1-180 again exhibited a high number of unique modified proteins (133), reinforcing observations from panel a.

**Figure 6:**
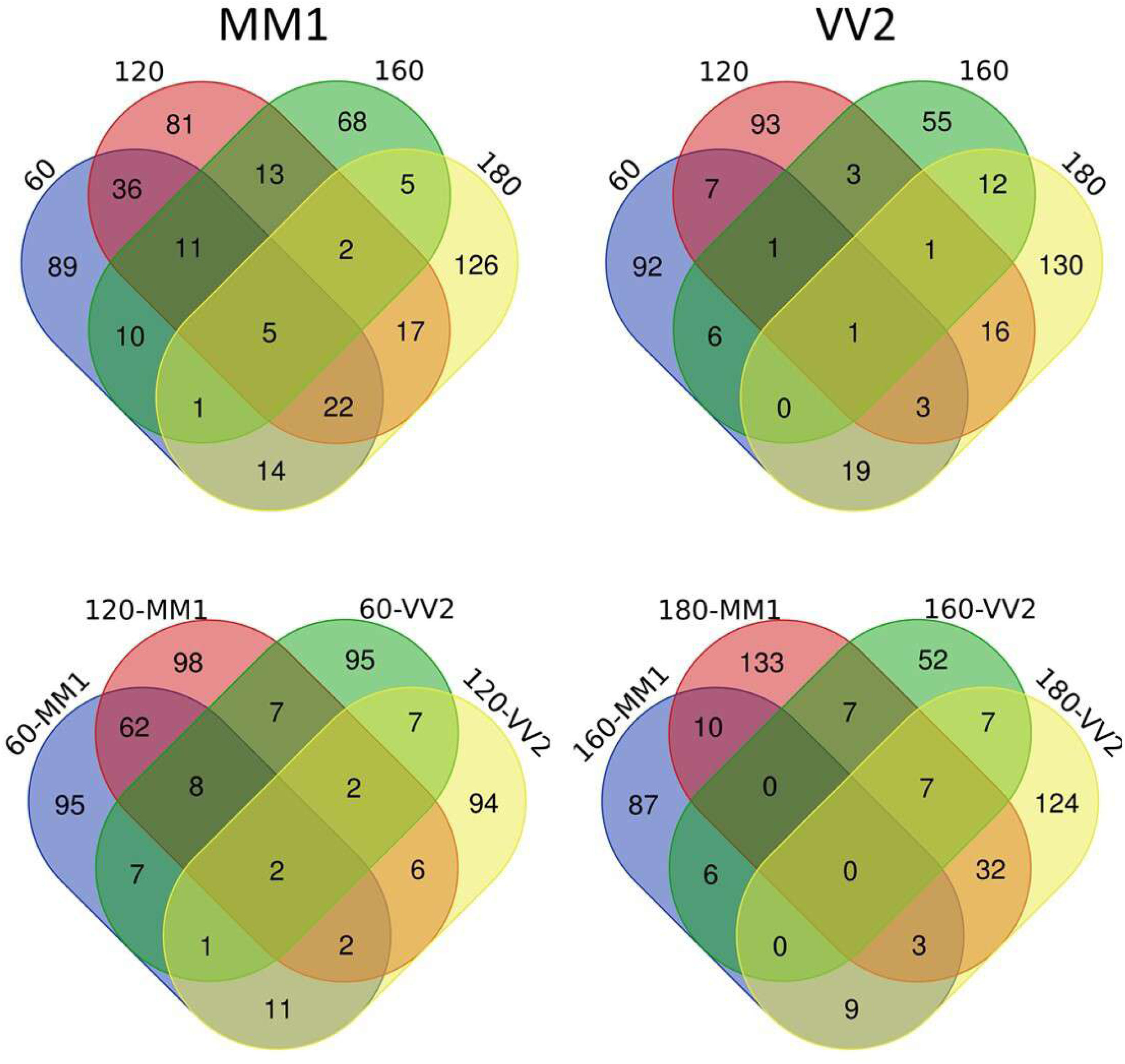
Shared and unique proteins between CJD-MM1- and CJD-VV2- inoculated mice. **A)** Common differentially expressed proteins at early preclinical (60 dpi), late preclinical (120 dpi), early clinical (160 dpi) and late clinical (180 dpi) time points, in (**a**) MM1-CJD-inoculated mice and (**b**) CJD-VV2-inoculated mice. **c)** Intergroup comparison between both subtypes at early preclinical (60 dpi), late preclinical (120 dpi), and **d**) at early clinical (160 dpi) and late clinical (180 dpi) time point. Dpi: days post inoculation.

### 3.3 Functional Profiling and Enrichment Analysis of Proteomic Targets in Prion Disease Progression

Next, to identify molecular events associated with the prion disease progression, we compared the functional profile of proteomic targets at each time point. For a detailed analysis, up-regulated and down-regulated proteins were analyzed separately. To perform functional enrichment analysis, we used ToppCluster (https://toppcluster.cchmc.org/), which allows the comparison of functional overrepresentations that differ as a function of time in time-series experiments. Currently, there are no existing tools that possess the capability to investigate multiple gene lists together and provide a functional modular map that includes a rich set of annotations. Networks generated using the program ToppCluster are displayed in Figure 7a and b, where major Gene Ontology (GO) annotations in terms of functional groups (Molecular Function: MF, Biological Process: BP, Cellular Component: CC, Pathway and Phenotype) are described (detailed annotation of the networks is provided in (Figure S5). As a confirmation, we also did a second functional enrichment analysis using WebGestalt GeneSet Enrichment Analysis (GSEA) (2).

**Figure 7:**
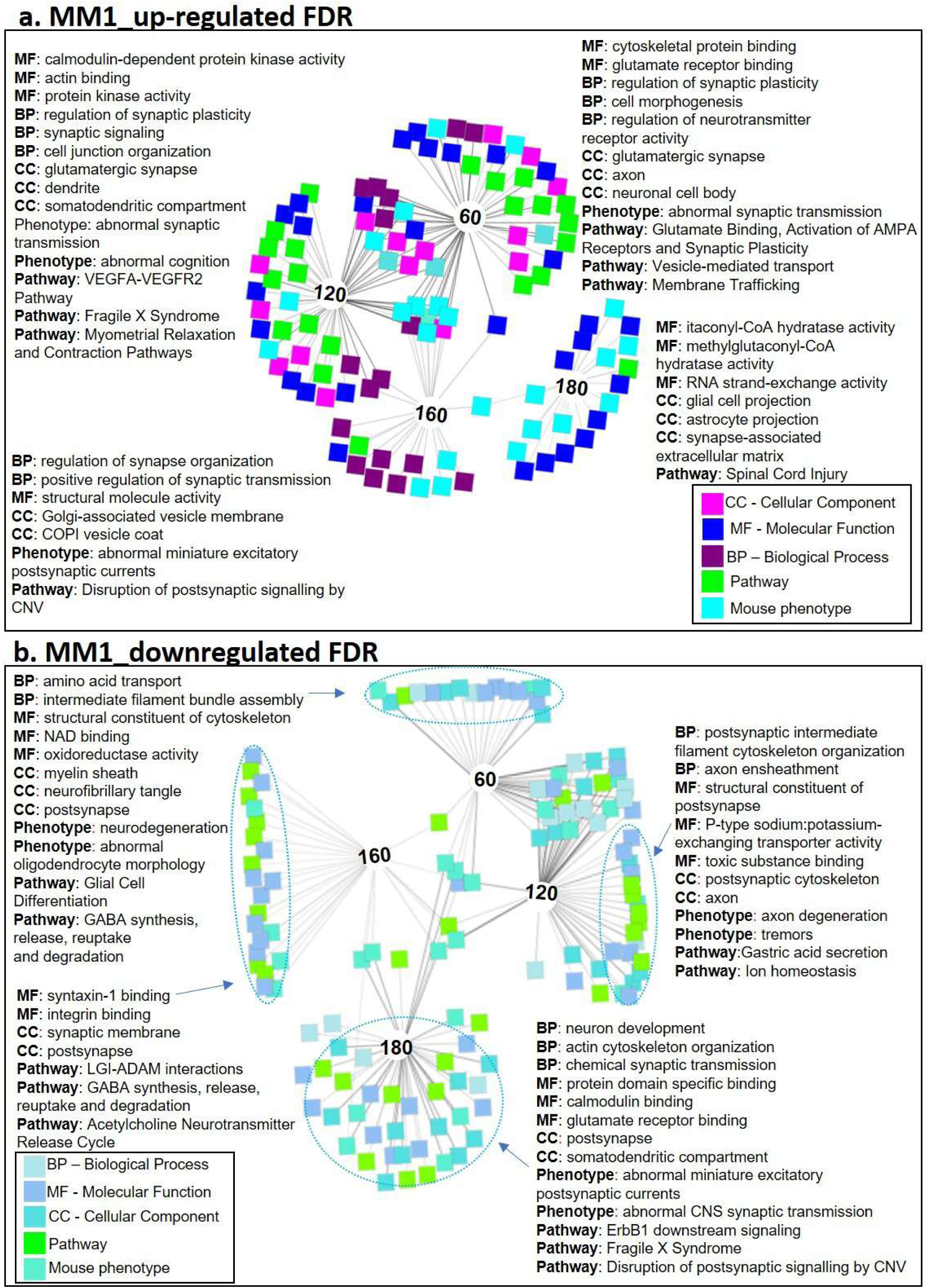
Functional profiling of significantly regulated proteins in CJD-MM1- inoculate mice. An ‘Abstracted’ Network showing enriched GO terms at early preclinical (60 dpi), late preclinical (120 dpi), early clinical (160 dpi) and late clinical (180 dpi) for **(a)** upregulated and **(b)** downregulated proteins using ToppCluster. BP: Biological Process, MF: molecular function, CC: cellular component, Mouse phenotype and Pathways. Dpi: days post inoculation.

Up-regulated proteins at the pre-clinical stage (60 dpi) showed significant enrichment for abnormal synaptic transmission. At late prelicinal stage (120 dpi), most significant functional terms were ‘’calmodulin-dependent protein kinase activity’’ and ‘’abnormal cognition’’ phenotype. At early preclinical (160 dpi), the most significant GO terms were ‘’regulation of synapse organization’’, structural molecule activity and ‘’abnormal miniature excitatory postsynaptic currents’’ as a phenotype. The most striking finding at the late clinical stage was enrichment of ‘’glial and astrocytic projections’’ and ‘’RNA strand-exchange activity and cellular component differences.

## 4. Discussion

The current study provides significant insights into the early stages of Creutzfeldt-Jakob Disease (CJD) infection, particularly regarding the MM1 and VV2 prion subtypes. By combining high-throughput proteomics and RT-QuIC assays, we were able to characterize the temporal progression of prion seeding activity and the molecular alterations associated with disease onset in humanized transgenic mice models. Our findings emphasize the importance of early biomarkers and proteomic changes in the context of CJD pathogenesis, which could serve as key indicators for early diagnosis and potentially provide targets for therapeutic intervention.

### 4.1 Early Detection of Prion Seeding Activity in Preclinical Stages of CJD

A key finding of our study is the detection of prion seeding activity in the brain well before the onset of clinical symptoms, highlighting the potential of early diagnostics in prion diseases. Utilizing the highly sensitive RT-QuIC assay (Orrú CD 2015), we identified detectable seeding activity as early as 60 days post-inoculation (dpi) in transgenic mice infected with human CJD-MM1 and CJD-VV2 prion strains. This early detection aligns with prior studies demonstrating preclinical prion accumulation in specific brain regions, such as the cortex and cerebellum (Zafar S 2015), and underscores the utility of RT-QuIC in detecting prions at asymptomatic stages.

Our data revealed notable differences in the seeding kinetics between the MM1 and VV2 models. MM1-infected mice showed a longer lag phase in RT-QuIC amplification, whereas VV2-infected mice exhibited minimal early-stage seeding but more robust responses at later time points. These strain-specific patterns are consistent with earlier findings on prion strain variability in propagation and disease onset (Peden AH 2004, Zafar S 2015). Furthermore, the spatial distribution of seeding activity—more pronounced in the cortex for MM1 and in the cerebellum for VV2—supports the concept of tissue-specific prion accumulation during disease progression (Zafar S 2018, Tarozzi M 2022).

Importantly, the progressive increase in RT-QuIC positivity from preclinical to clinical stages mirrors the temporal dynamics of prion replication (Candelise N 2017, Zerr I, 2020) and reinforces the potential of RT-QuIC as a non-invasive, early diagnostic tool. The observed strain-specific lag times also open the door to subtype-specific diagnostic strategies, offering a path toward more precise clinical assessments of prion disease subtypes.

### 4.2 Proteomic Alterations and Early Biomarkers of Disease

Proteomic analysis through SWATH-MS revealed profound alterations in the protein profiles of prion-infected mice, even at early preclinical stages. Notably, the proteins identified, such as Pdk1, Vps51, Lmnb2, Ahsa1 in CJD-MM1 and Dock3, Tnc, Ca8 and Gm10358 in CJD-VV2, are involved in critical cellular processes, including cell signaling, endocytosis, mitochondrial dysfunction, and cytoskeletal organization. These alterations at the early stages of disease suggest that prion replication triggers significant shifts in cellular machinery long before the onset of clinical symptoms, supporting the hypothesis that early molecular changes are a direct consequence of prion accumulation (Moreno et al. 2013, Kim 2020).

Furthermore, the temporal progression of proteomic changes was strikingly distinct between CJD-MM1 and CJD-VV2 models. The identification of distinct protein clusters at early preclinical stages for both prion strains (Fig. 7 and 8) suggests that each prion subtype induces a unique proteomic response, with potential implications for understanding strain-specific neuropathogenesis. For instance, alterations in synaptic transmission and cognitive function-related pathways were found at early preclinical stages in both prion models, while astrocytic and RNA processing pathways became more pronounced at later clinical stages (Fig. 7 and 8). These findings are consistent with other studies demonstrating that prion diseases lead to synaptic dysfunction and gliosis early in the disease process (Cunningham C 2005, Davies MJ 2015, Slota JA 2022). Moreover, the upregulation of synaptic proteins at early stages may suggest synaptic dysfunction as an early manifestation of prion disease, which may provide a window for intervention before irreversible damage occurs.

**Figure 8:**
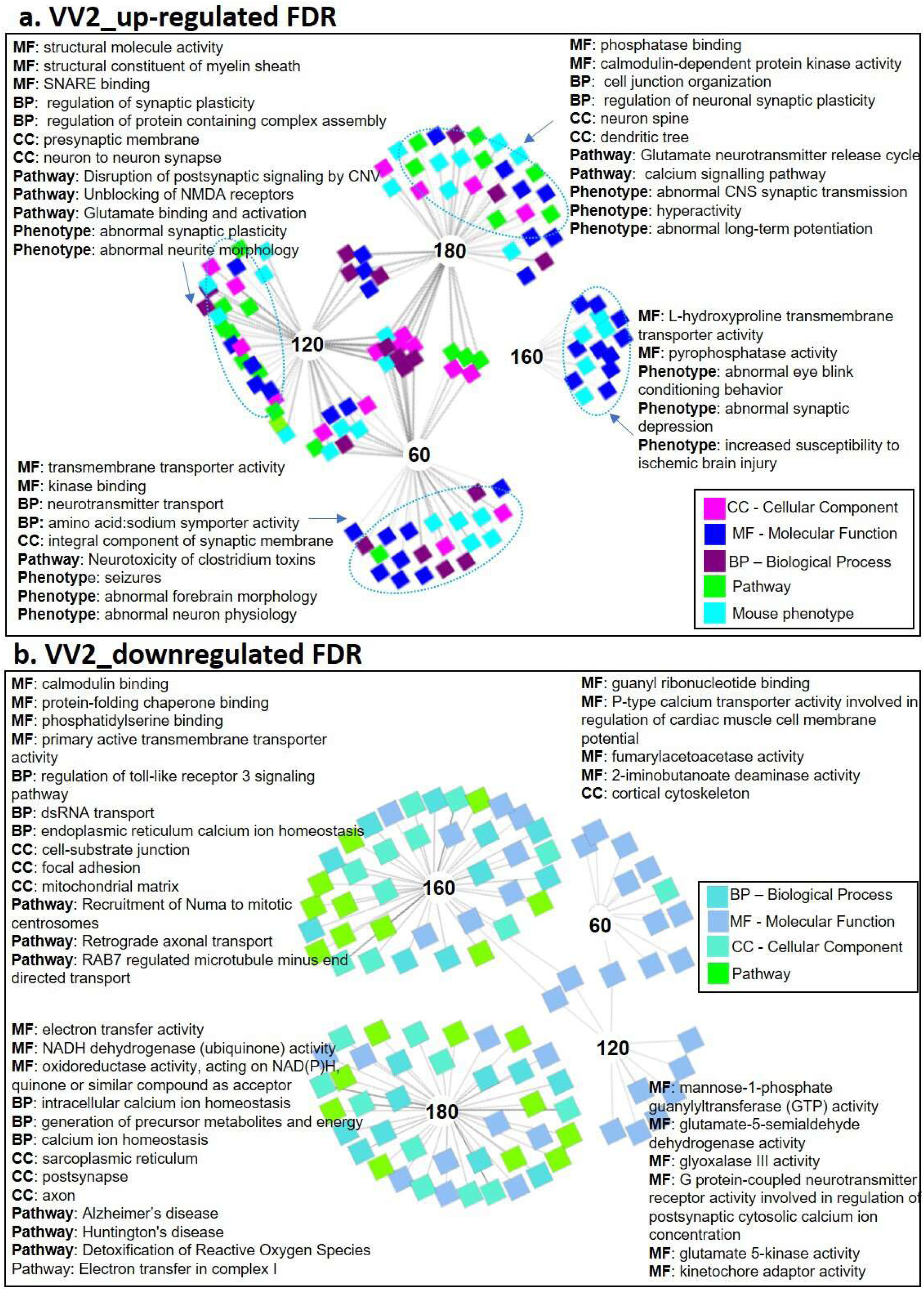
Functional profiling of significantly regulated proteins in CJD-VV2- inoculate mice. An ‘Abstracted’ Network showing top ten enriched GO terms at early preclinical (60 dpi), late preclinical (120 dpi), early clinical (160 dpi) and late clinical (180 dpi) for **(a)** upregulated and **(b)** downregulated proteins using ToppCluster. BP: Biological Process, MF: molecular function, CC: cellular component, Mouse phenotype and Pathways. Dpi: days post inoculation.

The proteomic analysis not only helps identify specific biomarkers associated with prion disease progression but also suggests possible therapeutic targets. For instance, proteins involved in synaptic plasticity, such as Pdk1, could be targeted to modulate synaptic function and potentially slow disease progression. Additionally, the differential expression of proteins in the two prion strains may offer insights into the molecular mechanisms underlying strain-specific pathogenesis, providing a basis for more tailored therapeutic approaches.

### 4.3 Functional Pathways and Implications for Disease Mechanisms

The functional enrichment analysis of differentially expressed proteins further emphasized the temporal changes occurring in prion-infected mice. At the early preclinical stage, synaptic transmission-related pathways were significantly enriched, pointing to early disruptions in neuronal communication as a key event in prion disease pathogenesis. As the disease progressed to later stages, there was a shift toward pathways related to astrocyte activity and RNA processing, suggesting that glial cells and non-coding RNA regulation become increasingly involved in the disease process and have already been reported in other neurodegenerative diseases (Zhao 2021, Nassar A 2022, D’Sa K 2025). This progression of molecular changes mirrors the clinical symptoms observed in prion diseases, where early cognitive and motor dysfunctions give way to more generalized neurological decline (Mallucci GR 2007, Harder A 2004).

We found disrupted cellular pathways in Creutzfeldt-Jakob disease (CJD), with the downregulation of genes implicated in ribosome biogenesis, synaptic transmission, cytoskeletal stability, vesicular trafficking, and proteostasis. GNL1 (Guanine Nucleotide-Binding Protein-Like 1), known for its role in ribosomal biogenesis and nucleolar function, is essential for maintaining protein synthesis and cellular homeostasis (Turcotte MA 2021). Its downregulation in CJD may compromise ribosomal integrity, leading to decreased translational capacity and reduced neuronal stress tolerance. This impairment aligns with evidence from ribosome profiling studies that reveal progressive translational dysfunction in glial cells during prion disease (Scheckel C 2020).

STXBP1 (Syntaxin-Binding Protein 1), critical for synaptic vesicle docking and neurotransmitter release, is another gene whose downregulation could significantly affect neuronal communication (de Wit H 2006, Toonen RF 2007, Südhof TC 2009, John A 2021). Loss-of-function mutations in STXBP1 are associated with severe neurodevelopmental disorders such as early-onset epileptic encephalopathies (Dilena R 2016), and reduced STXBP1 levels in CJD may reflect synaptic pathology that underpins the characteristic neurophysiological decline observed in prion disease (JA. 2010). Impairment of vesicle trafficking and neurotransmission could thus exacerbate synaptic degeneration, one of the earliest events in prion-related neurodegeneration.

PLLP (Plasmolipin), which encodes a proteolipid involved in the formation of the myelin sheath and neuronal membrane organization, is also significantly downregulated. Myelin abnormalities, including vacuolation and demyelination, have been described in CJD subtypes (Matsusue E 2004, Caverzasi E 2014), and reduced PLLP expression could contribute to destabilized axonal insulation and altered membrane dynamics. Such changes may increase neuronal susceptibility to excitotoxic and oxidative insults.

GPS1 (G Protein Pathway Suppressor 1), a negative regulator of the JNK signaling cascade, is implicated in modulating apoptosis and cellular stress responses (Li JY 2007). Its downregulation may relieve inhibition of the JNK pathway, leading to enhanced pro-apoptotic signaling and promoting neuronal death—a feature widely observed in neurodegenerative conditions (Xia Z 1995, Hunot S 2004). This may be particularly relevant in the context of prion diseases, where oxidative stress and neuroinflammation play prominent roles (Younas N 2023, Sun Z 2025).

The observed downregulation of NEFH (Neurofilament Heavy Chain), a key component of the neuronal cytoskeleton, further supports a model of structural degeneration. NEFH is crucial for maintaining axonal caliber and facilitating axonal transport (Zhao 2021), and its reduced expression may indicate cytoskeletal destabilization, contributing to axonal degeneration and synaptic loss. Notably, increased levels of neurofilament light chain (NfL), a related protein, are established biomarkers for axonal injury in prion diseases (Zerr I 2021, Schmitz M 2022, Vallabh SM 2023).

We also observed reduced expression of AHSA1 (Activator of Hsp90 ATPase Activity 1) functions as a co-chaperone within the protein quality control system, assisting Hsp90 in folding and stabilizing client proteins (Biebl MM 2019, Liu X 2022, Mondol T 2023). In prion diseases, characterized by progressive misfolding and accumulation of prion protein (PrP^Sc^), reduced AHSA1 expression could impair proteostasis mechanisms, exacerbating proteotoxic stress and facilitating prion propagation (Chernova TA 2017, Thellung S 2022, McKinnon C 2016). This aligns with broader models of neurodegeneration wherein failure of chaperone-mediated folding enhances the neurotoxic burden.

RALA (RAS Like Proto-Oncogene A), a small GTPase involved in vesicular trafficking and cytoskeletal organization, is essential for neuronal maintenance and communication (Lalli G 2005). Reduced RALA expression may disrupt intracellular trafficking, leading to impaired synaptic vesicle dynamics and accumulation of misfolded proteins, both of which are hallmarks of prion pathophysiology (Puig B 2016, Foliaki ST 2024). Similarly, CACYBP (Calcyclin Binding Protein) downregulation may disrupt calcium homeostasis and the ubiquitin-proteasome system, both vital to neuronal function and survival (Topolska-Woś AM 2016). Given the known link between calcium dysregulation and neurotoxicity (Gil-Martins E 2025), and the role of CACYBP in protein degradation (Fukushima T 2006, Qian F 2021), its suppression could enhance cellular stress and promote prion aggregation.

The upregulation of key regulatory proteins in MM1 Creutzfeldt-Jakob Disease (CJD) mice at the early pre-clinical stage provides valuable insight into the cellular changes that precede overt neurodegeneration. These molecular alterations highlight early disruptions in metabolic signaling, synaptic regulation, vesicular trafficking, and stress responses, which may serve as initial pathological hallmarks of prion disease.

One of the most prominent changes observed was the upregulation of pyruvate dehydrogenase kinase 1 (Pdk1), which plays a central role in shifting cellular metabolism from oxidative phosphorylation toward aerobic glycolysis by inhibiting the pyruvate dehydrogenase complex (Zhang et al., 2014). This metabolic reprogramming is a common neuroprotective strategy during early neurodegeneration, potentially aimed at reducing mitochondrial-derived reactive oxygen species (ROS) and oxidative stress. However, this shift may also lead to energy inefficiency and may not be sustainable as the disease progresses.

In brief, early pre-clinical MM1 CJD mice, the upregulation of genes such as Pdk1, Unc13a, Rab21, and Rraga reflects early metabolic shifts, synaptic changes, and increased vesicle trafficking—hallmarks of early neuronal stress. Pdk1 suggests a move toward glycolysis to counter oxidative stress (Zhang 2014), while Unc13a and Ppp1r9a indicate compensatory synaptic remodeling (Augustin 1999). Rab21 and Vps51 upregulation suggests enhanced endosomal-lysosomal activity to manage misfolded PrP (Zafar S, 2017, Liu P 2022, Luo L 2011). Eif4b and H2afx point to increased protein synthesis and DNA damage response (Chen B 2021, Mah 2010), and Atp2b2 reflects early calcium dysregulation (Smits JJ, Consortium und Koole W 2019). These changes suggest that neurons initiate stress responses well before clinical signs, offering potential early biomarkers and therapeutic targets.

In early-stage VV2 CJD mice, downregulation of Calb2, Hpcal1, and Dnaja1 suggests early deficits in calcium signaling, synaptic buffering, and chaperone-mediated proteostasis (Qiu XB 2006, Spilker C 2003, Coronas-Samano G 2016). Loss of Tnc, an extracellular matrix glycoprotein, may impair glial-neuronal communication and structural integrity (Tucić M 2021). However, reduced Ppm1h hints at disrupted phosphatase control in neuroinflammation (Nazish I 2021).

Conversely, upregulation of Prkca and Dock3 reflects heightened activity in synaptic plasticity and axonal remodeling (Wang H 2025, Makihara S 2018), while Syn2 and Agap2 suggest compensatory mechanisms for synaptic vesicle cycling and cytoskeletal stabilization (Medrihan 2013, Chouinard FC 2022). Mitochondrial genes like Ndufa4 and Tmem126a indicate metabolic adaptations (Xu LL 2025), and Fdps upregulation may signal altered isoprenoid biosynthesis and oxidative stress responses (Wang S 2022). Collectively, these changes highlight early disruptions in calcium homeostasis, mitochondrial metabolism, and synaptic signaling as potential drivers of VV2-specific prion pathology.

Overall, at the early pre-clinical stage, MM1 and VV2 CJD mice exhibit distinct molecular signatures indicative of subtype-specific neuronal stress responses. MM1 mice upregulate genes linked to glycolysis (Pdk1), synaptic remodeling (Unc13a), and vesicular trafficking (Rab21), alongside markers of DNA repair (Eif4b, H2afx) and calcium regulation (Atp2b2). In contrast, VV2 mice show early deficits in calcium buffering and proteostasis (downregulation of Calb2, Hpcal1, Dnaja1) and extracellular matrix integrity (Tnc). Compensatory responses in VV2 include upregulation of genes involved in synaptic plasticity and cytoskeletal dynamics (Prkca, Dock3, Syn2), as well as mitochondrial and metabolic adaptations (Ndufa4, Fdps). While both subtypes activate early compensatory pathways, MM1 emphasizes metabolic reprogramming and vesicle clearance, whereas VV2 shows greater disruption in calcium signaling and structural stability, suggesting divergent early pathogenic mechanisms with potential implications for subtype-specific biomarkers and interventions.

Collectively, these transcriptional/translational alterations reveal a broad-based failure of neuronal maintenance mechanisms in Creutzfeldt–Jakob disease (CJD). Rather than isolated defects, the convergence of dysregulated gene expression across critical pathways including protein synthesis, cytoskeletal integrity, synaptic function, and proteostasis reflects a systemic collapse of neuronal homeostasis. Whether these changes represent primary pathogenic drivers or downstream responses to prion accumulation remains to be clarified. Importantly, distinct CJD subtypes such as MM1 and VV2 exhibit differential gene regulation patterns, emphasizing the need for subtype-specific investigations.

Notably, both CJD-MM1 and CJD-VV2 models demonstrate similar functional pathway disruptions during early preclinical stages, though with varying intensities. This strain-dependent variation underscores the importance of understanding prion strain-specific mechanisms to inform the development of more precise diagnostic and therapeutic strategies. As prion diseases often share overlapping clinical features, these molecular distinctions at the protein and pathway levels may prove critical for improved subtype differentiation in clinical practice, ultimately enhancing patient stratification and enabling targeted therapeutic interventions. Deciphering the functional impact of these transcriptional changes could thus provide valuable insights into disease mechanisms and identify novel avenues to restore disrupted neuronal processes in prion and related neurodegenerative diseases.

## 5. Conclusions and Future Directions

In conclusion, our study underscores the power of combining RT-QuIC and high-throughput proteomics to gain a deeper understanding of prion diseases, particularly in the early preclinical stages of infection. The early detection of prion seeding and the identification of distinct proteomic alterations in MM1 and VV2 models suggest that these approaches can reveal critical biomarkers and molecular events that precede clinical symptoms. Moving forward, these findings could have significant implications for improving early diagnosis, tracking disease progression, and developing targeted therapies for prion diseases.

Future studies will need to extend these findings to human patient samples to validate the biomarkers identified here, particularly in asymptomatic individuals at risk for prion disease. Additionally, investigating the molecular mechanisms driving the differential seeding activity and protein alterations observed between prion strains could provide crucial insights into the strain-specific nature of prion diseases. Ultimately, a better understanding of these early molecular events may offer new opportunities for therapeutic intervention at the earliest stages of prion disease.

## Figures legends

**Supplementary Table S1.**

List of significantly altered proteins identified at preclinical and clinical time points in MM1-CJD. Proteins were identified by quantitative proteomics and filtered based on significance (adjusted p < 0.05). The table includes fold changes, time points of alteration (60, 120, 160, and 180 dpi), and functional annotations. Early-stage changes (60 dpi) mark the onset of molecular alterations coinciding with prion seeding activity detected by RT-QuIC.

**Supplementary Table S2.**

List of significantly altered proteins identified at preclinical and clinical time points in VV2-CJD. Differentially expressed proteins were determined as in Table S1. Distinct temporal profiles were observed, with early changes aligning with the initial detection of prion seeding. The table includes fold changes, significance values, and predicted functional roles.

